# Assembly processes inferred from eDNA surveys of a pond metacommunity are consistent with known species ecologies

**DOI:** 10.1101/2023.12.12.571176

**Authors:** Wang Cai, Maximilian Pichler, Jeremy Biggs, Pascale Nicolet, Naomi Ewald, Richard A. Griffiths, Alex Bush, Mathew A. Leibold, Florian Hartig, Douglas W. Yu

**Affiliations:** State Key Laboratory of Genetic Resources and Evolution, Kunming Institute of Zoology, Chinese Academy of Sciences, Kunming, China; Key Laboratory of Tropical Forest Ecology, Xishuangbanna Tropical Botanical Garden, Chinese Academy of Sciences, Mengla 666303, China; Theoretical Ecology, University of Regensburg, Regensburg, Germany; Freshwater Habitats Trust, First Floor, Bury Knowle House, North Place, Old High Street, Headington, Oxford, UK; Oxford Brookes University, Headington, Oxford, OX3 0BP, UK; Durrell Institute of Conservation and Ecology, School of Anthropology and Conservation, University of Kent, UK; Lancaster Environment Centre, Lancaster University, UK; Department of Integrative Biology, University of Texas at Austin, Austin, TX USA; Center for Excellence in Animal Evolution and Genetics, Chinese Academy of Sciences, Kunming, China; Yunnan Key Laboratory of Biodiversity and Ecological Security of Gaoligong Mountain, Kunming Institute of Zoology, Chinese Academy of Sciences, Kunming, China; School of Biological Sciences, University of East Anglia, Norwich Research Park, Norwich, UK

**Keywords:** Aquatic eDNA, metabarcoding, novel community data, biodiversity, pond, joint species distribution model (JSDM), macroecology, *Triturus cristatus*

## Abstract

Technological progress is enabling ecologists to create repeated, large-scale, structured, and standardised community surveys. However, it is unclear how best to extract information from these novel community data. We metabarcoded 48 vertebrate species from their eDNA in 320 ponds in England and applied the ‘internal-structure’ approach, which uses joint species distribution models to explain community compositions as the outcome of four metacommunity assembly processes: environmental filtering, dispersal, species interactions, and stochasticity. We find that the environment plays an important role in community assembly and that the inferred environmental preferences of species are consistent with their ecologies. We also infer negative biotic covariances between fish and amphibians, which is consistent with predator-prey interactions, and high spatial autocorrelation for the palmate newt, which is consistent with its hypothesised relictual distribution. Comparing sites in the metacommunity, environmentally and spatially distinctive sites are better explained by their environmental covariates and geographic locations, respectively, revealing sites where environmental filtering and dispersal limitation act more strongly. Furthermore, species belonging to different trait groups differ in how well environmental covariates, biotic covariances, and geographical locations explain their distributions. Overall, our results highlight the value of a modern interpretation of metacommunity ecology that embraces the fact that assembly processes differ between individual species and sites. We discuss how novel community data make feasible several study-design improvements that will strengthen the inference of metacommunity assembly processes from observational data.

## Introduction

Metacommunity theory, which explicitly models feedback between local communities and regional species pools, has been proposed as a unifying theory of spatial community ecology (Leibold *et al*. 2004; Leibold & Chase 2018). In the framework of this theory, we consider a set of communities whose local population dynamics are governed by environmental filtering, species interactions, and ecological drift and that are additionally linked by dispersal. The goal of the theory is to understand how these four basic assembly processes determine species compositions in the metacommunity (Vellend 2016). Traditionally, this has been done with classical community data gathered by human observers, but the fact that modern sensors such as eDNA deliver community observations that are ideally suited for metacommunity analysis has created excitement in the field and also makes metacommunity analysis interesting for molecular ecologists (Hartig *et al*. 2024).

Empirical approaches to studying metacommunities mainly aim at inferring the relative contributions of the four assembly processes (dispersal, environmental filtering, species interactions, and drift) from empirical data. Examples of these are analyses based on community summary statistics, such as ordinations that describe different metacommunities using centroids and distances, and alpha and beta diversities (Figure 1). Another common approach to analyse metacommunity data is variation partitioning, where classically, community composition is explained by metacommunity-level contributions of environmental and spatial factors (e.g. Cottenie 2005). However, those approaches exhibit limited power to reveal assembly processes (Guzman *et al*. 2022; Ovaskainen *et al*. 2019), in part because summary metrics cannot reveal how the four processes differentially affect individual species and sites. Leibold *et al*. (2022) refer to such metacommunity-level metrics as studying the ‘external structure’ of metacommunities, because they assume that there is one “average” mix of assembly processes that is the same across sites and species.

**Figure 1.**
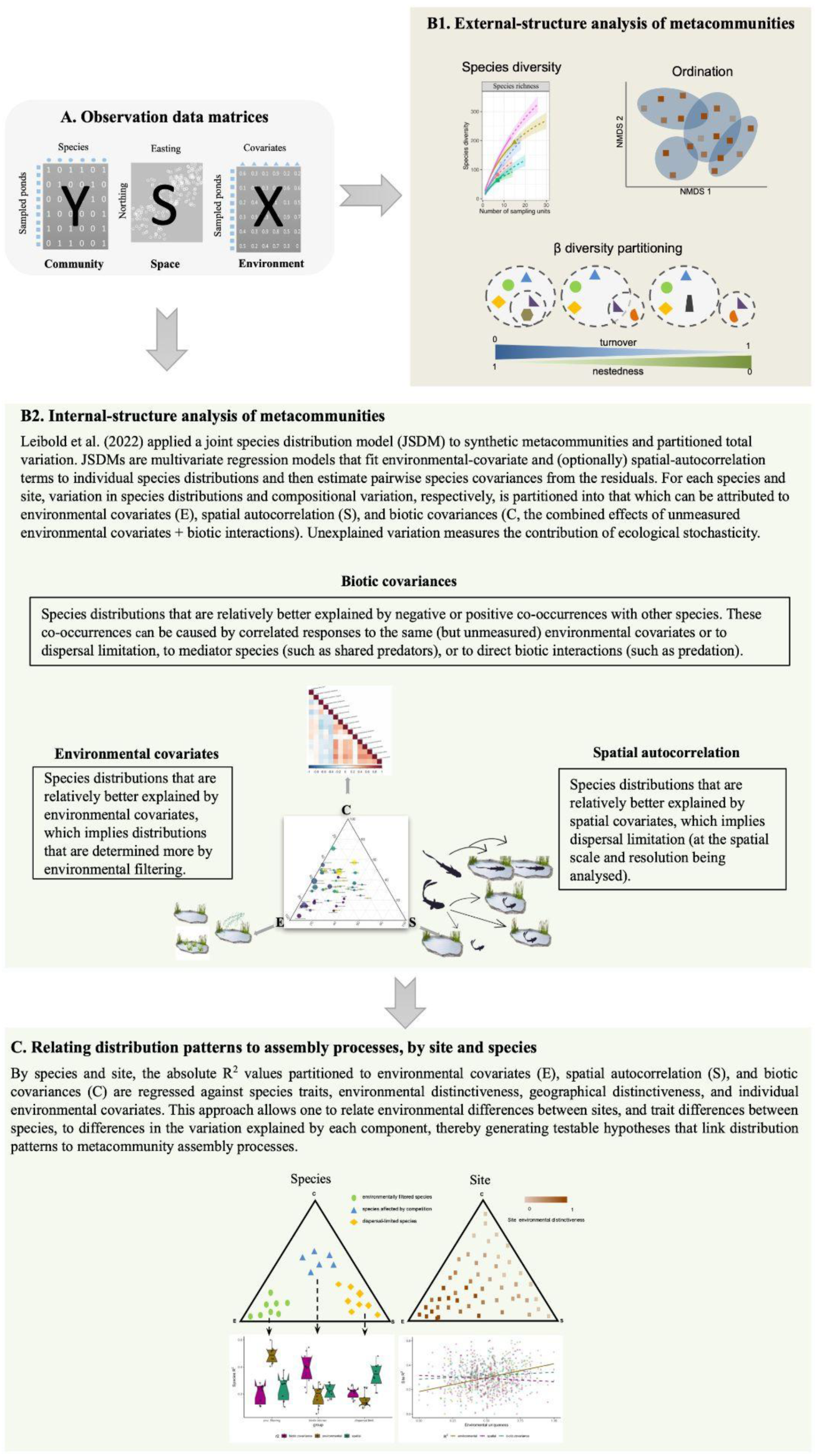
External-versus internal-structure analysis of metacommunities.

To avoid averaging assembly processes across sites and species, Leibold *et al*. (2022) propose to study the ‘internal structure’ of metacommunities, which dissects the importance of different assembly processes *by each species and site*. Technically, this can be done by using a joint species distribution model (JSDM) to partition the varying contributions of three model components (environmental covariates, species covariances, and spatial autocorrelation) to explaining species presence/absence, for each individual species and site (Figure 1). Among other things, *this approach allows one to relate environmental differences between sites, and trait differences between species, to differences in the variation explained by each component*, thereby generating testable hypotheses that link distribution patterns to metacommunity assembly processes.

Simulation studies have shown that internal-structure analysis can indeed differentiate synthetic metacommunities that differ in site environmental distinctiveness and in species niche breadth, dispersal ability, niche centrality, and the presence or absence of competitive interactions (Figure 1, Leibold *et al*. 2022; Terry *et al*. 2023). While these simulation results are encouraging, real metacommunity datasets have more complicated properties, including detection failures, measurement errors, and model uncertainty; not all species, environmental covariates, and sites can be included; species interact in multiple ways; and real metacommunities might be non-stationary, not least because of climate change (Abrego *et al*. 2021; Kadoya *et al*. 2024; Terry *et al*. 2023). Thus, it is important to gain more experience about the applicability of the internal structure idea to real data.

An ideal empirical metacommunity dataset for inferring internal structure would consist of (1) many local-community inventories with standardised species presence-absence or abundance information, (2) over an area that is connected (and large) enough for dispersal (and dispersal limitation) to operate, (3) with the taxonomic breadth to include interacting guilds such as predators and prey, and (4) measures of local environmental conditions relevant to the niche requirements of all these species. An exemplary study is provided by Kadoya et al. (2024) who applied internal-structure analysis to gillnet survey data covering three countries, 93 fish species, and 1,853 lakes and found that environmental covariates explained the most variation in species distributions and lake compositions, highlighting the importance of environmental filtering. Kadoya et al. then projected the effect of future climate heating on lake species compositions by running the fitted model with higher values of the degree-days environmental covariate while using the biotic covariances to simulate the effect of species interactions.

A promising alternative to traditional community observations is eDNA metabarcoding, which can generate repeated, large-scale, structured, and standardised community surveys (Hartig *et al*. 2024), but eDNA has rarely been used in metacommunity ecology. To our knowledge, the only attempt is by Vass *et al*. (2022), who metabarcoded aquatic fungi from four separate inlets along the Swedish coast and found in an internal-structure analysis that most species and site variance was attributable to environmental covariates, again highlighting the importance of environmental filtering. Vass *et al*. also found high biotic covariance values for some species, which is consistent with additional contributions from species interactions or unmeasured environmental covariates (Blanchet *et al*. 2020; Dormann *et al*. 2018; Hartig *et al*. 2024; Poggiato *et al*. 2021; Zurell *et al*. 2018).

Here, we test how well the internal structure of a candidate pond metacommunity surveyed with eDNA matches expectations derived from external knowledge of species ecologies. Our survey data come from ponds in the South Midlands of England that were originally sampled to detect the great crested newt (*Triturus cristatus*), a UK-protected amphibian species that breeds in ponds (Biggs *et al*. 2015). We metabarcoded the residual eDNA to detect vertebrates, generating a community matrix of 320 ponds `X` 48 vertebrate species. Each pond was associated with 8 environmental covariates and a geographic location, allowing us to fit three data matrices in a JSDM (Figure 1).

Among other results, we report at the species level that the fitted environmental covariates are consistent with known fish and amphibian ecologies, that the negative biotic covariances between fish and amphibians are consistent with expected predator-prey relationship between the two taxa, and that the high spatial autocorrelation for the palmate newt is consistent with a hypothesised relictual distribution for this species. At the site level, environmentally distinctive ponds are better explained by their environmental covariates, and spatially distinctive ponds are better explained by their geographic locations, revealing sites where environmental filtering and dispersal limitation act more strongly, respectively.

## Methods

### Environmental covariates

To quantify land cover around each pond, we used Rowland *et al*.’s (2017) 21 UK land classes. For each pond, we calculated the proportions of land class within a 500 m radius of its point location and used principal component analysis in {FactoMineR} v.2.4 (Lê *et al*. 2008) to extract the top three principal components (accounting for 40% of total variation, Figure S1), which correlate with the degree of agriculture vs. urban cover, grassland cover, and woodland cover. Each pond was also scored during sampling for ten standard pond variables used by surveyors to calculate the pond’s Habitat Suitability Index (HSI) for the great crested newt (ARG-UK 2010), of which we used five (Table S1).

### Pond sampling and newt assays

The pond water samples were the result of a single-season, great crested newt survey of 544 ponds in the South Midlands of England, UK in 2017, covering ponds in urban/suburban, woodland and agricultural areas. The survey used a grid design to generate single-species presence-absence data as input to a species distribution model that underpins a biodiversity offset market (Box 3 in Hartig *et al*. 2024). Samples were collected and processed following Biggs *et al*. (2015) and were stored at ambient temperature until shipped to a commercial lab (NatureMetrics, Egham, UK), and DNA was extracted using a precipitation protocol (Tréguier *et al*. 2014), after which each sample’s DNA was cleaned and subjected to 12 separate qPCR tests. After the qPCR assays, the residual eDNA was stored at -80°C. We loosely estimate that the cost of sample collection and great crested newt assays, including effort getting permission to sample private properties, exceeded US$120,000. Pond eDNA collected for great crested newts thus has several advantages for metacommunity studies in that we have a standardised, professionally collected, commercially processed, and spatially contiguous set of eDNA samples, with standardised environmental covariates.

### Metabarcoding assays

In 2019, the residual eDNA samples were retrieved and subjected to metabarcoding at NatureMetrics (PCR) and at Kunming Institute of Zoology (library prep), using a two-step protocol targeting a 73-110 bp fragment of 12S ribosomal RNA (Riaz *et al*. 2011). In the first step, 12 replicates per sample were amplified and then pooled to maximise the targeted sequence (Lahoz-Monfort *et al*. 2016). In the second step, three independent PCR replicates were performed on the pooled samples using twin-tagging (Yang *et al*. 2021; Zepeda-Mendoza *et al*. 2016). The three independent (separately twin-tagged) PCR replicates per sample in the second step were pooled into three approximately equimolar libraries for bead purification, library preparation, and sequencing on an Illumina HiSeq platform (PE150) at Novogene Tianjin, China.

We processed the raw sequence data with the modified DAMe bioinformatics pipeline of Cai *et al*. (2021), where we filtered out PCR and sequencing errors by retaining only sequences present in ≥2 PCR replicates with a minimum copy number of 25 per PCR. After sequence clustering, we generated a table of 540 ponds by 74 OTUs (operational taxonomic units). We assigned taxonomies to the OTUs using PROTAX (Axtner *et al*. 2019; Somervuo *et al*. 2017), setting prior probabilities to 0.90 for a list of expected UK vertebrate species (Harper *et al*. 2018).

### Joint species distribution modelling

To fit JSDMs to the observed community data, we converted OTU read counts to presence-absence data. We retained only OTUs present in ≥5 ponds and only sites with ≥1 targeted OTU (= vertebrate species present in the United Kingdom), which reduced the number of OTUs from 74 to 48 and the number of ponds from 540 to 320. We assigned species-level taxonomies to OTUs that received ≥98% PROTAX probability of species assignment, and we classified the OTUs into 6 trait groups: fish, amphibians, perching birds, waterfowl, mammals, and domestic species. Domestic species are whose distributions we deemed as determined largely by humans (Table S2).

We fit our data with two distinct JSDM structures. One model was fit to all the species in our dataset, terrestrial and aquatic (320 ponds `X` 48 species), and the other model fit to only the aquatic species (amphibians and fish, 279 ponds `X` 15 species; fewer ponds because we excluded those without aquatic species). *A priori*, aquatic species should be more likely to be filtered by pond characteristics, which make up 5 of our 8 environmental covariates. Thus, we expect the aquatic species, especially the fish, to act more like narrow-niche species (Figure 1C, top row), and the terrestrial species to act more like broad-niche species (Figure 1C, bottom row).

All models were fitted using {sjSDM} v.1.0.6 (Pichler & Hartig 2021) running under R 4.2.2 (R Core Team 2022). We used a binomial likelihood and a multivariate probit link, linear main effects for the eight environmental covariates, and a DNN (deep neural net) spatial model (trend surface model). To avoid overfitting, a light elastic net regularisation (Zou & Hastie 2005) was applied to all regression slopes and weights of the DNN (model fitting details in Appendix S1).

### Internal structure of the metacommunity

After fitting the two models (aquatic, terrestrial + aquatic), we used the ANOVA functions implemented in sjSDM (based on Leibold *et al*. 2022) to partition the variation of each species’ distribution and each site’s composition across environmental covariates (E), spatial autocorrelation (S), and biotic covariances (C), without the shared fractions of the pairwise components. The results are visualised using ternary plots, where the positions of the species and sites reveal the relative contributions of E, S, and C: the metacommunity’s internal structure (Figure 1B2).

Leibold *et al*. (2022) found that environmentally more distinctive sites (at the ends of their one niche axis) received higher E values (implying a greater contribution of environmental filtering). To test for this result in a natural metacommunity, we regressed the individual pond absolute E, S, and C R^2^ values against pond environmental distinctiveness using quantile regression (50% quantile) (Fasiolo *et al*. 2021). We also tested the parallel hypothesis that geographically distinctive sites would have higher absolute S R^2^ values (implying a greater contribution of dispersal limitation). In both cases, we defined distinctiveness as the leading eigenvector of the corresponding environmental or geographical Euclidean distance matrix. We note that if predictor variables are collinear, bivariate correlations can be spurious and partial correlations calculated using multiple regressions should be preferred. However, environmental and geographic distinctiveness showed practically no collinearity (Figure S2).

### Model generality

Our results are the outputs of a complex model that includes a linear environmental structure with eight environmental covariates and a DNN spatial structure with 30 x 2 layers. Complex models run a risk of overfitting, so to estimate the risk of overfitting after elastic-net regularisation, we carried out a 20-fold cross-validation test with stratified multi-label sampling (Gunopulos *et al*. 2011; Szymański & Kajdanowicz 2017). In each of 20 rounds, 19 folds (304 sites) were used as a training dataset, and the fitted model was used to predict species compositions in the hold-out fold (16 sites). Model fitting used the same structure, regularisation strengths, and scaled environmental and spatial covariates as the original full model. For each round and species, we calculated from the training dataset an explanatory- performance AUC (Area Under the Curve) metric. We also used each hold-out fold to calculate a predictive-performance AUC for each species: a measure of model generality. The final explanatory and predictive AUCs per species are the means over 20 folds. Species with higher predictive AUCs are those whose fitted models are more general (details in Appendix S1).

## Results

### Internal structure of the pond metacommunity

When analysing aquatic species only, we find that the fish are bimodally arrayed along the E-C axis, with four species relatively better explained by environmental covariates (higher E values), and six species relatively better explained by biotic covariances (higher C values), which reflect the effects of unknown environmental covariates plus possible species interactions (Figure 2A).

**Figure 2.**
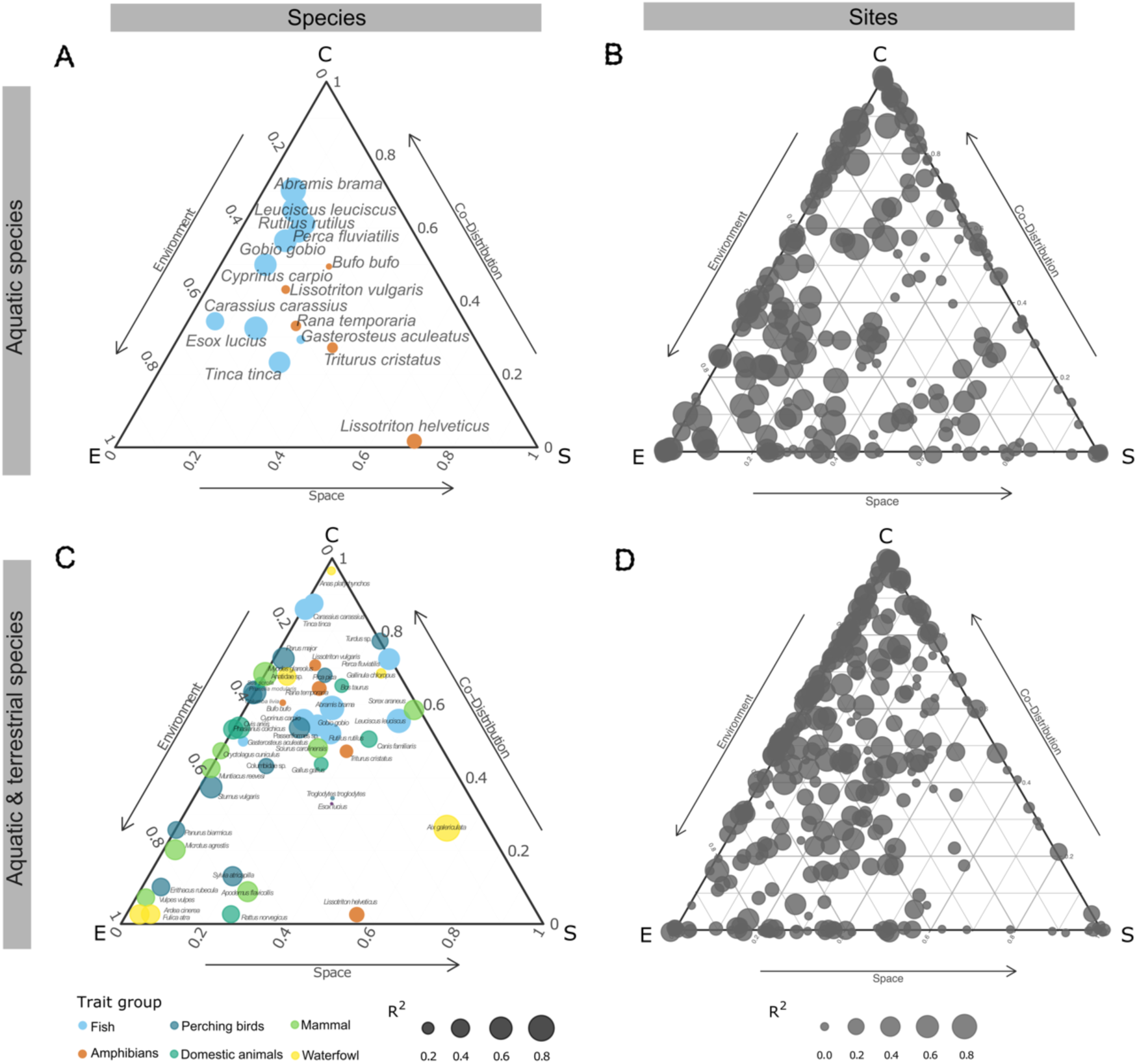
The internal structure of a pond metacommunity. The explained variation in species distributions or site compositions is decomposed, attributed to the three model components, environmental covariates (E), spatial autocorrelation (S), and biotic covariances (also known as co- distribution) (C), and visualised in a ternary plot after dividing each component’s explained variance by its sum to allow comparison among species. Top row. Aquatic species only. Bottom row. Aquatic + Terrestrial species. Left column. Each point is a species, point size scales to total R^2^_McFadden_ of each species, and the colours code for species trait group. Right column. Each point is a site (pond), and point size scales to total R^2^_McFadden_ of each site.

In contrast, none of the five amphibian species shows a high contribution of either environmental covariates or biotic covariances, but relative to fishes, amphibians show greater contributions of dispersal limitation (higher S values), especially the palmate newt (*Lissotriton helveticus*), whose distribution is mostly explained by dispersal limitation. This species was detected in 16 ponds, in three separate sections of the survey area (Figure S3), and its high S value indicates that other ponds in which this species was not detected have environmental conditions similar to those in which the palmate newt was detected.

Including terrestrial species in the analysis (Figure 2C) increases the relative contribution of biotic covariance for both fish and amphibians, which could reflect either the contributions of species interactions with terrestrial species or more unmeasured environmental covariates that have been revealed by adding the terrestrial species. The terrestrial species themselves also largely range along the E-C axis, with no clear clustering by trait group. Like the aquatic species, only one terrestrial species, Mandarin duck (*Aix galericulata*), has a high S value (Figure S3) and is found in only five ponds. For the two site ternary plots (Figure 2B, D), the general effect of adding terrestrial species is an increase in the variance accounted for by biotic covariances (site points shift upwards toward C).

We now examine species and site variation to try to infer some of the assembly processes that have resulted in these observed internal structures.

### Estimated environmental preferences

In the aquatic-only model (Figure 3A), the pond effects for the fish species are in the direction of greater prevalence in *larger* ponds with *lower* risk of drying and *less* macrophyte cover. Several of the fish species are known to eat macrophytes, reduce macrophyte cover through other behaviours, and/or require higher oxygen with less macrophyte cover (Lopes *et al*. 2015; Maceda-Veiga *et al*. 2017; Stefanoudis *et al*. 2017). In contrast, for the amphibians, pond effects are in the direction of greater prevalence in *smaller* ponds with *higher* macrophyte cover. Pond drying risk, water quality, and shade showed essentially no effects on amphibian prevalence. Most of the effects of land cover on fish species are in the direction of lower prevalence in areas surrounded by agriculture or grassland. Grassland here is dominated by the ‘improved grassland’ subtype (Rowland *et al*. 2017), characterised by regular fertilisation, herbicide application, and drainage (DEFRA & Rural Payments Agency 2022). For amphibians, the only effects are in the direction of greater prevalence of palmate newt and common frog in ponds bordered by woodland. All five amphibian species may utilise woodland outside the breeding season (Mulkeen *et al*. 2017; Oldham *et al*. 2000).

**Figure 3.**
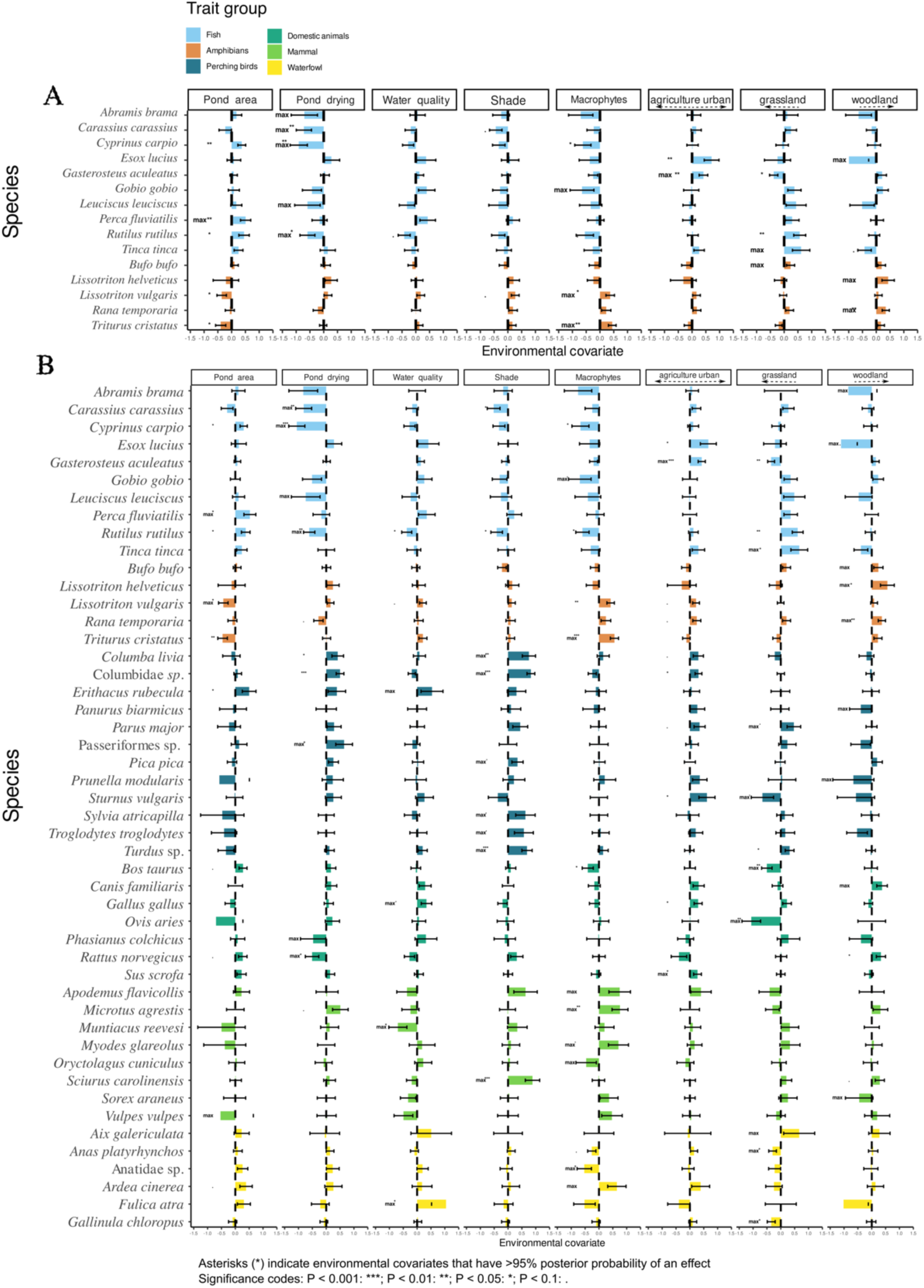
Estimated environmental effects. Eight environmental covariates were included in the model. The first five covariates from the left are taken from the ten standard pond variables used by surveyors to calculate the Habitat Suitability Index (HSI) of each pond for the great crested newt and are therefore measured at all ponds in our dataset (ARG-UK 2010). The last three covariates describe the dominant land cover class within 500 m of each pond (more details in Table S1). Horizontal bars show the magnitudes, directions, and standard errors of the coefficients of each of the eight environmental covariates for each species. All covariates were normalised before fitting. Significance values are not corrected for multiple comparisons. Colours indicate species trait groups.

In the aquatic+terrestrial model (Figure 3B), the effects of the environmental covariates on fish and amphibians remain largely the same as in the aquatic-only model. For the terrestrial species, most of the significant environmental-covariate effects are shade (% of pond perimeter shaded by trees), macrophyte cover, and land cover. Shade, which affects many perching birds and the grey squirrel, is most parsimoniously interpreted as increasing species detectabilities. Macrophyte cover is positively correlated with three rodent species and the grey heron and negatively correlated with rabbits and ducks (Anatidae sp.). Land-cover effects are variable across species, but we note that cows and sheep have higher prevalences in ponds bordered by (‘improved’) grassland.

Finally, we observe that the Eurasian coot (*Fulica atra*) has higher prevalences at ponds of higher water quality, which in the HSI scoring system is inferred from the presence of an “abundant and diverse invertebrate community” (ARG-UK 2010).

### Biotic covariances

We visualise the residual biotic covariances in pairwise correlation plots (Figure 4), where large absolute correlation-coefficient values correlate with high C-values in the internal-structure ternary plots (Figure 2A and 2C; linear model, aquatic species only, R^2^ = 0.726, P < 0.001; aquatic + terrestrial, R^2^ = 0.201, P < 0.001).

**Figure 4.**
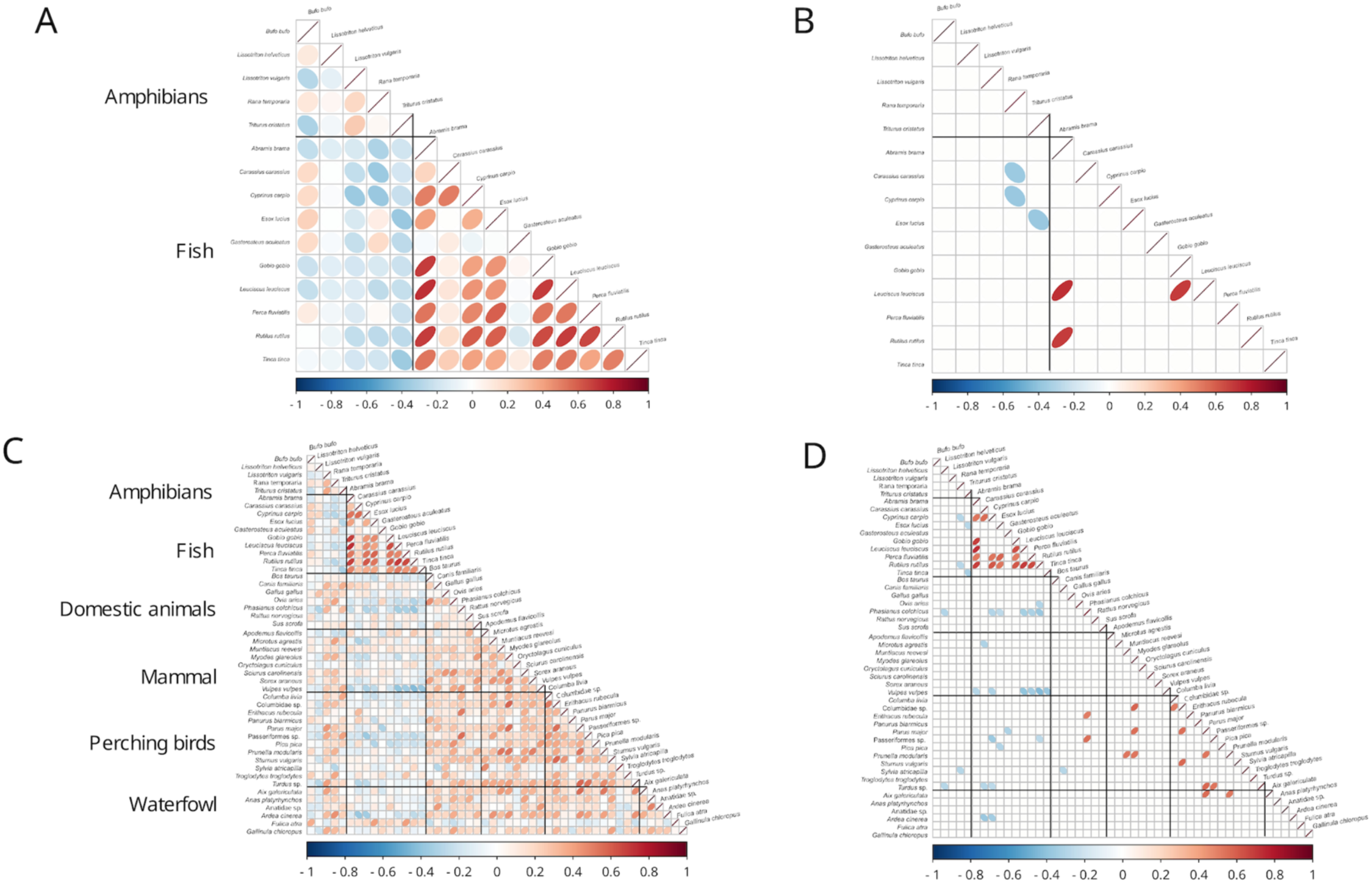
Biotic covariances. A. Aquatic species only, all pairwise covariances. B. Covariances filtered to the 2.5% most negative and positive. C. Aquatic + terrestrial species, all pairwise biotic covariances. D. Covariances filtered to the 2.5% most negative and positive.

In the aquatic species model (Figure 4A, B) and after filtering to the 2.5% most negative and positive values, the three surviving negative correlations are between the common frog (*Rana temporaria*) and two omnivorous fish species (*Carassius carassius* and *Cyprinus carpio*) and between great crested newt (*Triturus cristatus*) and a carnivorous fish (*Esox lucius*). There are also three surviving positive correlations between fish species, which we conservatively interpret as indicating unmeasured environmental covariates.

In the aquatic+terrestrial species model (Figure 4C, D) and after filtering to the 2.5% most negative and positive values, there are four surviving negative correlations between amphibians and fish. The common frog is negatively correlated with two omnivorous fish species (*Cyprinus carpio* and *Rutilus rutilus*), and great crested newt (*Triturus cristatus*) is negatively correlated with two carnivorous/omnivorous fish (*Esox lucius* and *Tinca tinca*). Most of the surviving positive correlations occur among the fish species and among the bird species, which we again interpret as unmeasured environmental covariates. Also notable are negative correlations between several fish species with ring-necked pheasant (*Phasianus colchicus*) and red fox (*Vulpes vulpes*).

### Relating distribution patterns to assembly processes, by site and species group

By site, the absolute R^2^ explained by the environment increases significantly with the environmental distinctiveness of the site (Figure 5A), and the absolute R^2^ explained by space increases significantly with the geographical distinctiveness of the site (Figure 5B). In other words, environmental filtering appears to be an increasingly more important assembly process for more environmentally distinctive sites, as predicted by Leibold et al. (2022), and dispersal limitation appears to be increasingly more important for geographically distinctive sites. Given this environment effect, we *post-hoc* tested each covariate individually and found that the absolute R^2^ explained by the environment increases only with pond area (Figures 5C, S4), suggesting that the species compositions of large ponds are determined more strongly by environmental filtering.

**Figure 5.**
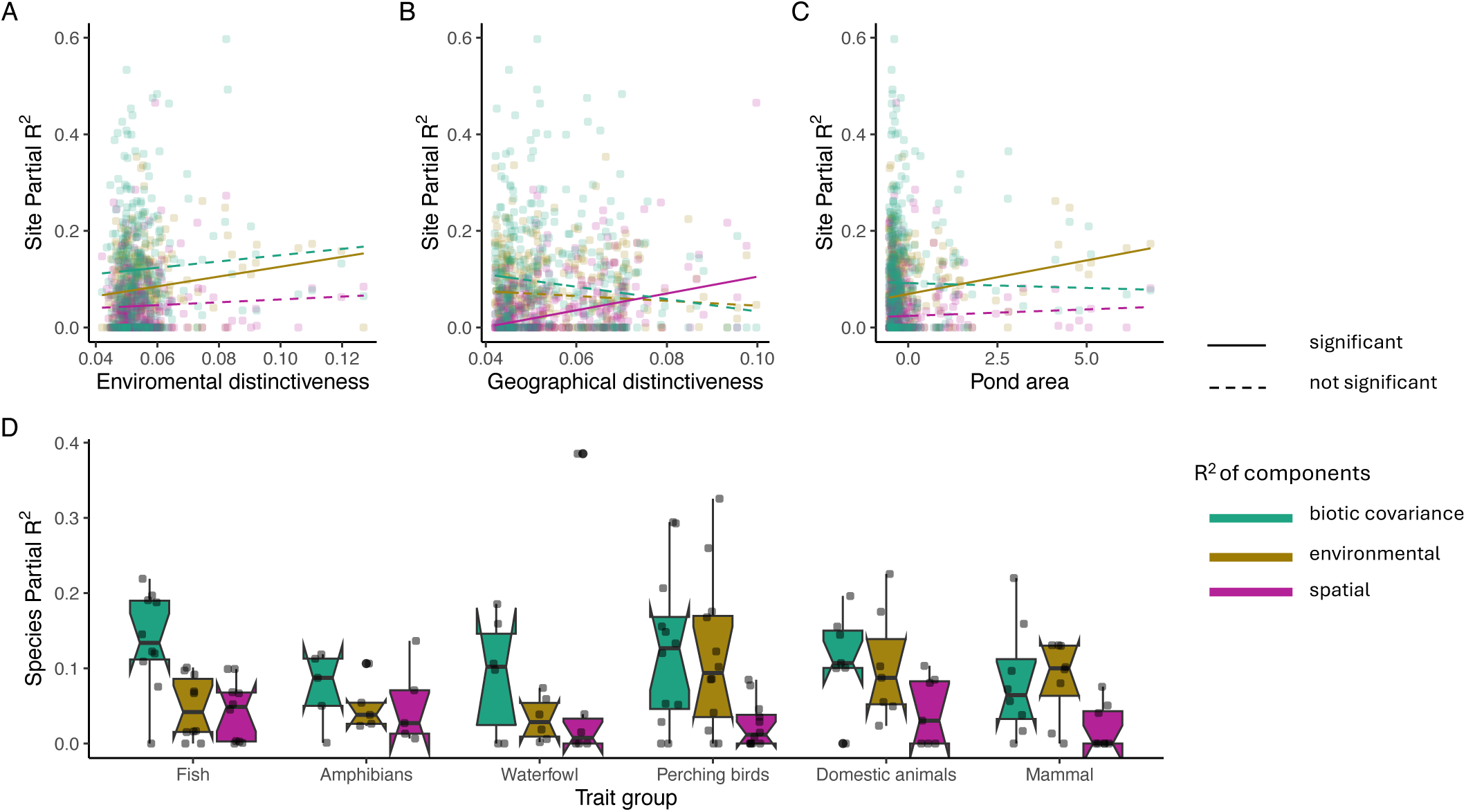
Correlation of the importance of assembly processes per site and species to environmental predictors and traits. A-C: Quantile regression, correlating the importance of the three assembly mechanisms (measured by the share of absolute partial R^2^ values, each in a different colour) per site against A. environmental distinctiveness. D. association of R^2^ shares per species with traits, in this case, species groups. Solid lines indicate significant regressions at P<0.05 significance level (p = 0.006 for the environmental component line), B. geographical distinctiveness (p < 0.001 for the spatial line), and C. pond area (p < 0.001 for the environmental line). Dashed lines are non-significant regressions. Note that the shared fractions were removed from the partial R^2^, so the three components will not necessarily sum up to the total R^2^ value, which is displayed in Fig. 2.

By species group, the absolute R^2^ explained by biotic covariances is greatest for fish, amphibians, and waterfowl, and about equal with the R^2^ explained by the environment for the other three trait groups (Figure 5D), which is consistent with species distributions being primarily governed by a combination of environmental filtering and (to a lesser extent) species interactions. Interestingly, the most pond-dependent species (fish, amphibians, waterfowl) are the least well explained by our environmental covariates, although there are individual exceptions (Figure 2A, C).

### Model generality

For the aquatic-only model, explanatory AUCs are always slightly greater than predictive AUCs, and explanatory and predictive AUCs are positively correlated (linear model, adjusted R^2^=0.546, P=0.001) (Figure 6A). For the aquatic+terrestrial model, explanatory AUCs are again still always greater than predictive AUCs, but the correlation weakens considerably (linear model, adjusted R^2^=0.139, P=0.005), and for some terrestrial species, the model makes worse-than-random predictions (predictive AUCs<0.5) (Figure 6B). The risk of overfitting is greater for low-predictive-AUC species, so the risk is greater for terrestrial species. Both models are generally better at predicting fish distributions than amphibian distributions. Palmate newt (*Lissotriton helveticus*) is an exception, with the overall second-highest predictive AUC (Figure 6A), which is probably explained by the large contribution of space to its explained variance. Stickleback (*Gasterosteus aculeatus*) and European perch (*Perca fluviatilis*) are exceptions for fish, with the overall two lowest predictive AUCs in the aquatic model (Figure 6A).

**Figure 6.**
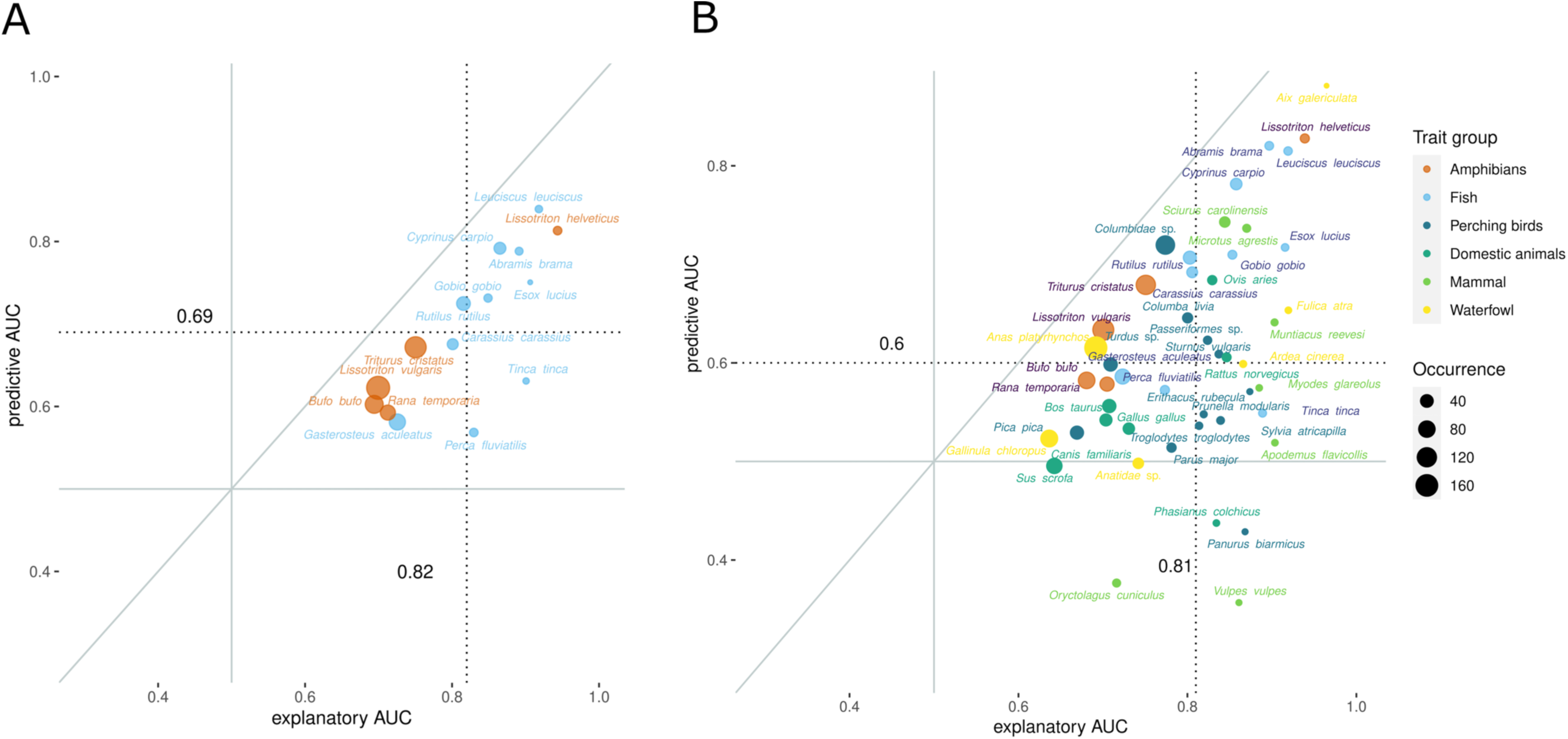
Predictive versus explanatory performance in two JSDMs. A. Aquatic-species-only model. B. Aquatic + Terrestrial species model. Model performance was assessed using the AUC metric, and the dotted lines indicate mean AUC values (vertical for predictive and horizontal for explanatory). In both models, predictive performance is generally higher for fish than for amphibians, and explanatory performance is generally somewhat greater than the predictive performance, indicating moderate overfitting.

## Discussion

The goals of our study were to reveal the internal structure of a real metacommunity, which infers the importance of different assembly processes per species and site. We used a joint species distribution model to estimate these relative and absolute contributions of environmental covariates, biotic covariances, and space for explaining spatial variation in pond compositions. Pondscapes are convenient study systems because (1) each pond is unambiguously identified as a local community, (2) there is an *a priori* division between likely pond niche specialists (aquatic species) versus generalists (terrestrial species), (3) aquatic eDNA metabarcoding can efficiently generate hundreds of local-community inventories, and (4) the detected species encompass multiple trophic levels, increasing the possibility of detecting species interactions.

### Importance of taxonomic breath

We inferred two internal structures, one for aquatic species only and one for aquatic + terrestrial species (Figure 2). This gives us our first conclusion, which is that the ‘one average mix’ approach to metacommunities indeed loses the useful information revealed in species and site variation (Leibold *et al*. 2022). In this pond metacommunity, even if one makes the extreme assumption that all the variation partitioned to biotic covariance is also environmental filtering (but unmeasured), the species and sites still vary in their inferred degrees of dispersal limitation, and thus no single metacommunity paradigm (i.e. species-sorting, mass-effect, patch-dynamic, and neutral communities; Holyoak *et al*. 2005; Shoemaker & Melbourne 2016; Suzuki & Economo 2021; Thompson *et al*. 2020) can serve as an adequate description.

### Influence of environmental effects

Looking at how metacommunity assembly processes vary across sites and species, both of the internal structures (Figure 2A, C) suggest that environmental filtering is an important structuring force for many of the species in this pondscape. These findings are supported by the fact that several of the environmental-covariate coefficient values are consistent with known biology. For instance, relative to amphibians, fish have a higher prevalence in larger, more permanent ponds (Figure 3). Macrophyte cover is also lower in ponds with higher prevalence of fish over amphibians, although the direction of causality might vary by species, since some fish species can directly reduce macrophyte cover. High macrophyte cover also appears to increase the prevalence of multiple terrestrial and waterfowl species (Figure 3) On the other hand, and contrary to its inclusion as a contributor to the suitability of habitats for newts (ARG-UK 2010), higher water quality is not associated with higher prevalence of any amphibian species, although it is associated with higher prevalences of Eurasian coot (Figure 3).

If environmental filtering is an important assembly process in this metacommunity, Leibold et al.’s (2022) corollary is that more environmentally distinctive sites should be more strongly determined by environmental filtering. This is what we observe (Figure 5A). In fact, this result holds separately for four of the six trait groups (Figure S5), the exceptions being fish, for which there is no relationship, and waterfowl, for which *spatial* site R^2^ increases with environmental distinctiveness.

### Influence on space on the community assembly

Looking at the contribution of space, which can be interpreted as a proxy for dispersal limitation, we find that spatial effects are dominant for two species, palmate newt and mandarin duck (Figure 2). The spatial distribution of the 16 palmate newt detections is visibly patchy (Figure S3), which may be due to the persistence of relictual populations with a historic distribution associated with woodland (Beebee & Griffiths 2000). There are only five detections for mandarin duck (Figure S3), so it is unclear whether such a spatial pattern implies dispersal limitation, but this introduced and free-living species is described as largely sedentary in the UK (Harris 2024).

In parallel to the environmental distinctiveness test, we also found that dispersal limitation appears to be stronger in geographically distinctive sites (Figure 5B).

### Influence of co-distribution on community assembly

With the main exception of common toad (*Bufo bufo*), which is protected from fish predation by toxins, residual associations between amphibians and fish are mostly negative, and three or four biotic correlations survive after filtering to the most extreme values (Figure 4). Keeping in mind the caveat that biotic covariances should not be taken as direct evidence for species interactions (Blanchet *et al*. 2020; Dormann *et al*. 2018; Hartig *et al*. 2024; Poggiato *et al*. 2021; Zurell *et al*. 2018), it is plausible that these associations arise from species interactions since the fish species involved are omnivorous or carnivorous and could thus prey on amphibian eggs, larvae, or possibly adults.

### What can eDNA bring to metacommunity ecology and internal structure analysis?

The large gains from eDNA metabarcoding in efficiency and error homogeneity over traditional survey methods make it feasible to generate datasets with many samples and many species, which can strengthen the inference of metacommunity assembly processes. Most obviously, the large number of species detectable with eDNA increases the probability of detecting sets of species that are truly interacting, as suggested by the negative correlations between predatory and omnivorous fish and amphibians from this study (Figure 4). Including more species could also reveal important but unmeasured environmental covariates via their pairwise species covariances. For instance, while terrestrial and aquatic species mostly do not interact directly, some terrestrial species could be proxies for unmeasured land uses that affect aquatic species, such as increased agricultural runoff. In our models, we included agricultural and improved-grassland land cover types as environmental covariates, so we saw this effect directly (Figure 3). The negative covariances of foxes and pheasants with multiple fish species (Figure 4D) might therefore be revealing other unmeasured land-use covariates (N.B. >35 million pheasants are released annually in the UK, mostly in England, and appear to boost fox numbers and incentivise land-cover management measures (Sage *et al*. 2020)).

More technically, eDNA sampling makes it more feasible to collect multiple sample replicates, which would allow combining a JSDM with a detection model to account for observation error (Diana *et al*. 2022; Doser et al. 2023; Guillera-Arroita *et al*. 2017; Hartig *et al*. 2024; Tobler *et al*. 2019). Also, only two species in our dataset showed strong signals of dispersal limitation (Figure 2), but this low number could be because near-neighbour ponds were not sampled in our dataset, removing the possibility of detecting fine-scale spatial autocorrelation and thereby possibly reducing the relative importance of dispersal that would support source-sink relations among closely adjacent ponds. Denser sampling might have detected more evidence of dispersal limitation. In our case, unfortunately, our dataset used the great crested newt sampling protocol, which requires only one sample per pond, and the ponds were dispersed across the landscape because the original use was to fit a species distribution model.

### What can JSDMs and internal structure analysis bring to eDNA researchers?

Novel community datasets, including eDNA metabarcoding, are multivariate response datasets; that is, each species is a response variable, and there are many of them. Before computing power was widely available, such datasets were generally first reduced to tractable dissimilarity matrices before visualisation and analysis (e.g. NMDS and constrained ordination), but this approach loses information and generates artefacts (Warton *et al*. 2012). However, for over a decade now, it has been possible to analyse multivariate abundance data directly (Warton 2022). For instance, we have shown here that JSDMs allow the simultaneous analysis of environmental covariates, biotic covariances, and spatial autocorrelations, and the outputs can be visualised and interrogated in powerful ways (Hartig *et al*. 2024; Popovic *et al*. 2019; Terry *et al*. 2023; van der Veen *et al*. 2022; Warton 2022).

## Conclusion

In conclusion, our study demonstrates that the combination of eDNA data and the analytical approach of exposing the ‘internal structure’ of metacommunities using JSDMs is a powerful tool for examining community assembly processes in structured landscapes. For the pond metacommunity in this study, we could reveal interesting patterns of how the importance of community assembly processes across sites and species, and relate those differences to spatial, environmental and trait predictors.

Overall, these results fall in line with other empirical studies (Kadoya *et al*. 2024; Mehner *et al*. 2021; Vass *et al*. 2022) that support Leibold et al.’s (2022) theoretical argument that JSDMs can reveal local (site and species-specific) assembly processes from static observational datasets, thereby making it possible to perform confirmatory tests of established ecological hypotheses or create new hypotheses from observational data. For instance, the *post-hoc* result that environmental site R^2^ increases with the pond area environmental covariate (Figure S4), suggests that environmental filtering is more important in larger ponds. This can in principle be tested by creating ponds of different sizes in the same general area and monitoring species colonisation.

Our study thus provides a blueprint for ecologists who want to study metacommunity assembly processes and also for molecular ecologists that want to extract more information from their eDNA data.

