## Supplementary Information for "Assembly processes inferred from eDNA surveys of a pond metacommunity are consistent with known species ecologies"

**The PDF file includes:**

- Supplementary methods
- Reference for supplementary Methods
- Table S1 and S2, and Figures S1 to S5

### Supplementary methods

***Pond sampling and GCN assays*** — Water samples were processed using standardised protocols from Natural England's GCN programme (Biggs *et al.* 2015). At each pond, water samples were collected evenly from 20 locations around the pond. 30 ml of water was collected from each of the 20 locations and then pooled to produce a 600 ml water sample for each pond. 90 ml of water was taken from the pooled sample to make six 15 ml subsamples. 33.5 ml absolute ethanol and 1.5 ml sodium acetate were added to these six subsamples and shipped at ambient temperature to NatureMetrics, Egham, UK. Samples were stored at -20°C until DNA extraction. During sampling, surveyors recorded ten metrics of pond quality which were used to calculate a single GCN Habitat Suitability Index (HSI, hereafter Oldham *et al.* 2000; ARG-UK 2010). In this study we used five of these as environmental variables (see Table 1). The deprecated metrics can be replaced by metabarcoding data and geographic location.

DNA extraction was performed according to the precipitation method (Tréguier *et al.* 2014), following the manufacturer's instructions for the DNeasy Blood & Tissue Extraction Kit (Qiagen GmbH, Hilden, Germany). The six subsamples from each pond were pooled into one, so that from now on there is only one extracted DNA sample for each pond.

After DNA extraction, the samples were first used to perform qPCR for the detection of great crested newt (*Triturus cristatus*) for Natural England's GCN programme in 2017. The qPCR assay was performed using TaqMan qPCR assays with the published primers and probe from Thomsen et al. (2012). Each sample was given a GCN 'eDNA score' from 0 to 12 (the number of positive qPCRs); any score >0 is legally treated as a positive detection (but see Diana *et al.* 2021). After qPCR assays, eDNA samples were stored at -80°C until metabarcoding analyses. ALL laboratory experiments were carried out in the commercial laboratory (NatureMetrics, Egham, UK) and after GCN assays, eDNA samples were donated to us for research.

***Metabarcoding protocol*** — In 2019, we reprocessed samples for vertebrate detection. All stored eDNA samples were first purified using the QIAquick PCR Purification Kit (Qiagen GmbH, Germany) prior to vertebrate amplification. PCR amplification was performed using a two-step protocol targeting a 73-110 bp fragment of 12S ribosomal RNA (Riaz *et al.* 2011). The PCR conditions were as follows: initial denaturation at 95 °C for 5 min, followed by 45 cycles (5 cycles for the second PCR step) of 95 °C for 15 s, 57 °C for 30 s, 72 °C for 30 s, and finishing at 72 °C for 5 mins. All PCRs were performed in 20 μL reactions containing 0.6 U Ex Taq HS DNA polymerase, 1 X Ex Taq buffer (Mg2+ plus), 0.2 mM dNTP mix, 0.4 μM of each primer, 1 μL DMSO, 0.1 μg/μL BSA and 2 μL eDNA sample. In the first step, 12 replicates per sample were amplified and then pooled to maximise the target sequence (Lahoz-Monfort *et al.* 2016). In the second step, three independent twin-tag PCR replicates were performed on the pooled samples following a modified DAMe metabarcoding protocol (Zepeda-Mendoza *et al.* 2016; Alberdi *et al.* 2018; Yang *et al.* 2021). This protocol uses different twin-tags (unique 7-9 nucleotide sequences added to both forward and reverse primers) for each of the three PCRs per sample, allowing them to be distinguished during bioinformatic processing. By taking advantage of the three PCR replicates with twin-tags, we can filter out numerous error sequences. Four PCR positive controls (mixed known fish sequences) and four negative controls (use of ultrapure water instead of DNA template) were included in each 96-well plate in both PCR steps to detect possible contamination. After the second PCR, three independent PCR replicates per sample were pooled into three approximately equimolar libraries for bead purification (Agencourt AMPure XP Kit, Beckman Coulter, Inc., USA), library preparation (NEXTflex Rapid DNA-Seq Kit for Illumina (Bioo Scientific Corp., Austin, USA)) and sequencing (performed on the Illumina HiSeq platform, PE150) at Novogene Tianjing, China.

***Bioinformatic protocol*** — Sequencing resulted in 280, 557, 173 paired-end reads. Prior to sample demultiplexing, the raw data were trimmed for residual Illumina adapters and low-quality ends using *AdapterRemoval* 2.2.0 (Schubert *et al.* 2016) and *Sickle* 1.33 (Joshi & Fass 2011), then denoised using the *Bayes-Hammer* module in *SPAdes* 3.10.1 (Nikolenko *et al.* 2013), and finally the read pairs were merged using *PandaSeq* 2.11 (Masella *et al.* 2012). In all cases, default parameters were used. Subsequently, *Begum* (Gopalakrishnan 2022) was implemented to demultiplex the sequence into samples and filter out tag jumps, errors and low-quality sequences.

After removing a very large number of tag-jumped reads (66, 874, 449), Begum filtering was applied (we retain sequences in ≥ 2 of the 3 PCRs per sample with ≥ 25 copies per PCR and length between 80 bp - 120 bp). The remaining sequences were clustered into 98% Similarity Operational Taxonomic Units (OTUs) using *SUMACLUST* 1.0.20 (Mercier *et al.* 2013), and then {LULU} v 0.1.0 with default parameters was applied to filter out erroneous OTUs (Frøslev *et al.* 2017).

Probabilistic taxonomic assignments were obtained from the PROTAX software (Somervuo *et al.* 2017), following the workflow recommended by Axtner et al. (2019). We assigned taxonomies to OTUs using a trained PROTAX model, with a list of expected UK vertebrate species (for which prior probabilities were set to 0.90) taken from Harper et al. (2018) to avoid artificially increasing regional diversity through incorrect assignments.

We noted that some OTUs representing domestic species were also retained, although they were probably caused by contamination, we cannot confirm whether they were present or caused by other reasons such as meat products. These OTUs are used in the statistics as 'domestic species'. We also removed OTUs present in samples that were unlikely to be present in UK ponds. As these eDNA samples were not collected and designed for this study at the outset, they were stored with the Arica samples for two years before metabarcoding. We also removed OUTs present in both samples and negative controls, which were matched to humans and species used as positive controls. Finally, 74 OUTs representing resident species in the UK regional pool were retained, leaving 351 samples with at least one OTU (351 sites X 74 OTUs). We assigned species-level taxonomies to OTUs that received ≥ 98% PROTAX probability of species assignment, otherwise OTUs would receive a genus or family name if the taxonomic assignment at that level was ≥ 98%.

To obtain higher quality data to fit the model, we further filter sites that lack environmental variables and location (7 sites were removed) and contain possible false positive OTUs (23 sites were removed). We assume that qPCR provides more accurate detection (Harper et al. 2018), so if metabarcoding detection of GCN shows positive detection but qPCR does not, we define these detections as false positives and remove these sites. This reduced the OTU table to 321 sites X 74 OTUs.

***Joint Species Distribution Modelling*** — We only keep species that had ≥5 occurrences, which means they are less likely to be false positive OTUs, 26 OTUs and one site were removed. This leaves 320 sites with 48 OTUs (species present in the United Kingdom) for the following analysis. 41 of the 48 OTUs are wild species and 7 OTUs are domestic species (Table S1). To fit a JSDM to the observed community data, we converted OTU read numbers to presence-absence (binary) data. We classified OTUs into 6 groups: fish, amphibians, perching birds, waterfowl, mammals, and domestic species. Domestic species are those whose distributions are determined by humans (Table S1).

To analyse the internal structure of the metacommunity, we fitted two joint species distribution models (jSDM) using the {sjSDM} package v.1.0.6 (Pichler & Hartig 2021). The first model was fitted to all the species in our dataset, terrestrial and aquatic (320 ponds X 48 species) and the second model only to the aquatic species (amphibians and fish, 279 ponds X 15 species, fewer ponds since we excluded those without aquatic species). Models were fit with binomial likelihood and multivariate probit link, linear main effects for all 8 environmental covariates, and a deep neural networks (DNN) spatial model. To avoid overfitting a light elastic net (Zou & Hastie 2005) regularisation was applied to all regression slopes and weights of the DNN. The model is run with a batch size of 0.1*sites, a learning rate of 0.01, and 50000 samples for Monte Carlo integration, an iteration control when the model is best fit, it stops the model to continue traning (after 150 iterations, reaching optimal fit), or finishes 500 iterations if no best fit is reached. The model was tuned with the light linear regularization strength (lambda.env = 0.001, alpha.env = 1.0) for environmental structure, the light linear regularization strength (lambda.bio = 0.001, alpha.bio = 1.0) for biotic covariance, and a DNN structure for spatial structure (two layer deep neural network, 30 neurons for each layer, with regularization strength of lambda.sp = 0.002, alpha.sp = 0.2). These hyperparameters also apply to prediction model.

***Model generality*** — Our results (Figures 2, 3, 4) are the outputs of a complex model that includes a linear environmental structure with eight environmental covariates and a deep neural network spatial structure with 30 x 2 layers. Complex models run a risk of overfitting, so to estimate the risk of overfitting after elastic-net regularisation, we carried out a 20-fold cross-validation test, using the *IterativeStratification* module of the *Scikit-multilearn* Python package, for stratified multi-label sampling, to partition the data (Gunopulos *et al.* 2011; Szymański & Kajdanowicz 2017). In each of 20 rounds, 19 folds (304 sites) were used as a training dataset, and the fitted model was used to predict species compositions in the hold-out fold (16 sites). Model fitting used the same structure, regularisation strengths, and scaled environmental and spatial covariates as the original full model, as described above. For each round and species, we calculated from the training dataset an explanatory-performance AUC (Area Under the Curve) metric, which equals 1 for a model with 100% correct predictions and 0 for 100% incorrect predictions. We also used each hold-out fold to calculate a predictive-performance AUC for each species: a measure of model generality. The final explanatory- and predictive-performance AUCs per species were the means over all 20 folds. Species with higher predictive AUCs are those whose fitted models are more general (Figures 2, 3, 4).

**Table S1. Environmental covariates included and excluded from the** JSDMs**.** The first ten covariates are used to calculate the Habitat Suitability Index (HSI) value of a pond for great crested newt (ARG-UK 2010). We omitted the last five, as they are redundant or potentially redundant with the metabarcoding data (waterfowl, fish scores, pond count), redundant with the land-cover covariates (terrestrial habitat) or lacked variation altogether (geographic location). The last three covariates are the first three principal components extracted from a PCA analysis of land cover within 500 m of each pond, with the land-cover classes extracted from Rowland et al. (2017).

| Environmental covariate | Included in model or not | Description |
|---|---|---|
| Pond area | Included in model | The surface area of the pond ($m^2$) when water is at its highest level (excluding flooding events) |
| Pond permanence | Included in model | How often a pond dries, from ‘never dries’ to ‘dries annually’. Ponds are scored by interviewing the landowner, and if this information is not available, the surveyor makes a judgement based on the water level at the time of the survey. For instance, a pond that is dry by late spring is likely to dry every year, so the pond is described as "Dries annually". |
| Water quality | Included in model | The rating is based on invertebrate diversity, good water supports an abundant and diverse invertebrate community, poor water quality supports low invertebrate diversity, and bad water quality is clearly polluted. |
| Shade | Included in model | The percentage of the estimated pond perimeter that is shaded to at least 1m from the bank, usually by trees. |
| Macrophytes | Included in model | The percentage of the estimated pond surface area occupied by macrophyte cover, including emergent, floating plants (excluding duckweed) and submerged plants reaching the surface. |
| Waterfowl | Not included | Description of waterfowl density where there is evidence of waterfowl presence and its effect on pond vegetation. We do not include this index because we used metabarcoding to detect waterfowl to species. |
| Fish | Not included | Presence of fish in the pond, based on local knowledge and surveyor observation. We do not include this in the model because we used metabarcoding to detect fish to species. |

| | | |
|---|---|---|
| Pond count | Not included | Number of ponds within 1 km of the survey pond and not separated by major dispersal barriers such as major roads. As samples were collected on a grid design, one per square kilometre, and surveyors only visit the sampled pond, we did not include this index. |
| Geographic location | Not included | Because all ponds in this dataset are located in ARG-UK's (2010) Geographical location Zone A, all ponds received the same score, so we do not include it. |
| Terrestrial habitat | Not included | Subjective surveyor rating of surrounding terrestrial habitat according to perceived suitability for great crested newt. As we are not only estimating great crested newt habitat, we do not include this covariate, but instead use the next three covariates to measure dominant local land cover classes. |
| Agriculture-urban | Included in model | The first principal component. Positive values indicate increasing arable land cover, and negative values indicate increasing urban land cover. For each pond, the cover of each land-cover class was counted from a satellite-derived land-cover map (Rowland *et al.* 2017). This 21-dimensional dataset was reduced to three principal components (Figure S1). |
| Grassland | Included in model | The second principal component. Negative values indicate increasing coverage of 'improved grassland'. |
| Woodland | Included in model | The third principal component. Negative values indicate increasing coverage of woodland. |

7

**Table S2. Species list and assigned trait group.** Incidence is the number of ponds in which each species was found.

| Trait group | Scientific name | Incidence | Trait group | Scientific name | Incidence | Trait group | Scientific name | Incidence |
|---|---|---|---|---|---|---|---|---|
| Amphibians | *Lissotriton vulgaris* (smooth newt) | 148 | Mammal | *Microtus agrestis* (field vole) | 11 | Perching birds | *Pica pica* (European magpie) | 43 |
| Amphibians | *Triturus cristatus* (great crested newt) | 121 | Mammal | *Oryctolagus cuniculus* (European rabbit) | 11 | Perching birds | *Columba livia* (pigeon) | 21 |
| Amphibians | *Bufo bufo* (common toad) | 82 | Mammal | *Muntiacus reevesi* (Reeve's muntjac deer) | 8 | Perching birds | *Parus major* (great tit) | 16 |
| Amphibians | *Rana temporaria* (common frog) | 54 | Mammal | *Apodemus flavicollis* (yellow-necked mouse) | 7 | Perching birds | Passeriformes sp. | 12 |
| Amphibians | *Lissotriton helveticus* (palmate newt) | 16 | Mammal | *Myodes glareolus* (bank vole) | 7 | Perching birds | *Sturnus vulgaris* (common starling) | 10 |
| Fish | *Gasterosteus aculeatus* (three-spined stickleback) | 64 | Mammal | *Sorex araneus* (common shrew) | 7 | Perching birds | *Sylvia atricapilla* (blackcap) | 10 |
| Fish | *Rutilus rutilus* (roach) | 43 | Mammal | *Vulpes vulpes* (red fox) | 6 | Perching birds | *Troglodytes troglodytes* (wren) | 9 |
| Fish | *Cyprinus carpio* (common carp) | 31 | Domestic species | *Sus scrofa* (pig) | 66 | Perching birds | *Erithacus rubecula* (European robin) | 8 |
| Fish | *Carassius carassius* (crucian carp) | 26 | Domestic species | *Gallus gallus* (chicken) | 45 | Perching birds | *Panurus biarmicus* (bearded tit) | 6 |
| Fish | *Perca fluviatilis* (perch) | 16 | Domestic species | *Bos taurus* (cow) | 34 | Perching birds | *Prunella modularis* (dunnock) | 6 |
| Fish | *Gobio gobio* (gudgeon) | 15 | Domestic species | *Canis familiaris* (dog) | 29 | Waterfowl | *Anas platyrhynchos* (mallard duck) | 167 |
| Fish | *Abramis brama* (common bream) | 14 | Domestic species | *Ovis aries* (sheep) | 16 | Waterfowl | *Gallinula chloropus* (common moorhen) | 84 |
| Fish | *Leuciscus leuciscus* (dace) | 13 | Domestic species | *Rattus norvegicus* (brown rat) | 15 | Waterfowl | Anatidae sp. (duck/geese/swan) sp. | 23 |
| Fish | *Tinca tinca* (tench) | 11 | Domestic species | *Phasianus colchicus* (common pheasant) | 7 | Waterfowl | *Ardea cinerea* (grey heron) | 8 |
| Fish | *Esox lucius* (northern pike) | 10 | Perching birds | Columbidae sp. (dove) sp. | 107 | Waterfowl | *Fulica atra* (coot) | 7 |
| Mammal | *Sciurus carolinensis* (grey squirrel) | 23 | Perching birds | *Turdus* sp. (thrush) sp. | 46 | Waterfowl | *Aix galericulata* (mandarin duck) | 5 |

**Figure S1. Principal component analysis (PCA) of land cover classes.** The PCA groups 21 land cover classes into ten dimensions. We initially included in the model the first five, all with eigenvalues greater than one, and ran a preliminary JSDM. The output revealed weak signals for PC4 and PC5, and we therefore used only PC1, PC2, and PC3 for the final model. A. PC1 and PC2. B. PC1 and PC3.

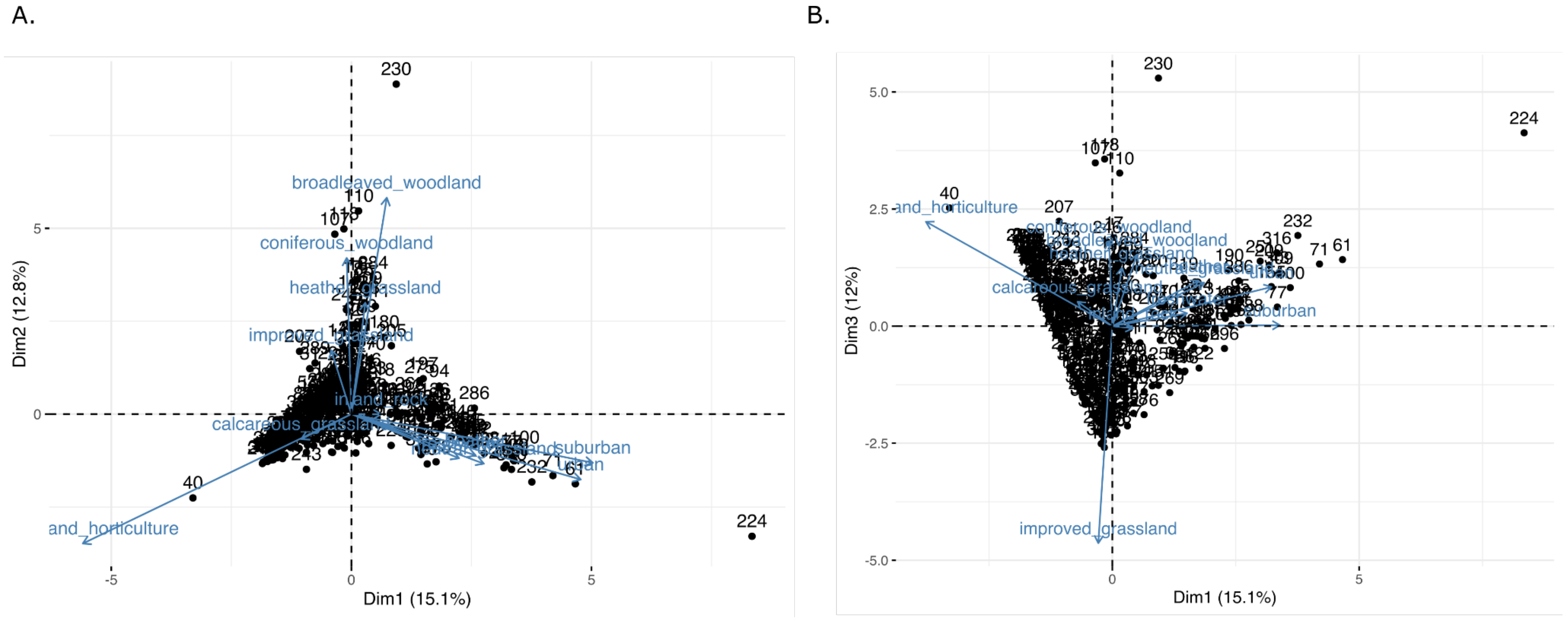

**Figure S2.** Pairwise correlations of the eight environmental covariates used in the model (Table S1), species richness, and environmental and spatial distinctiveness. Correlations are generally low.

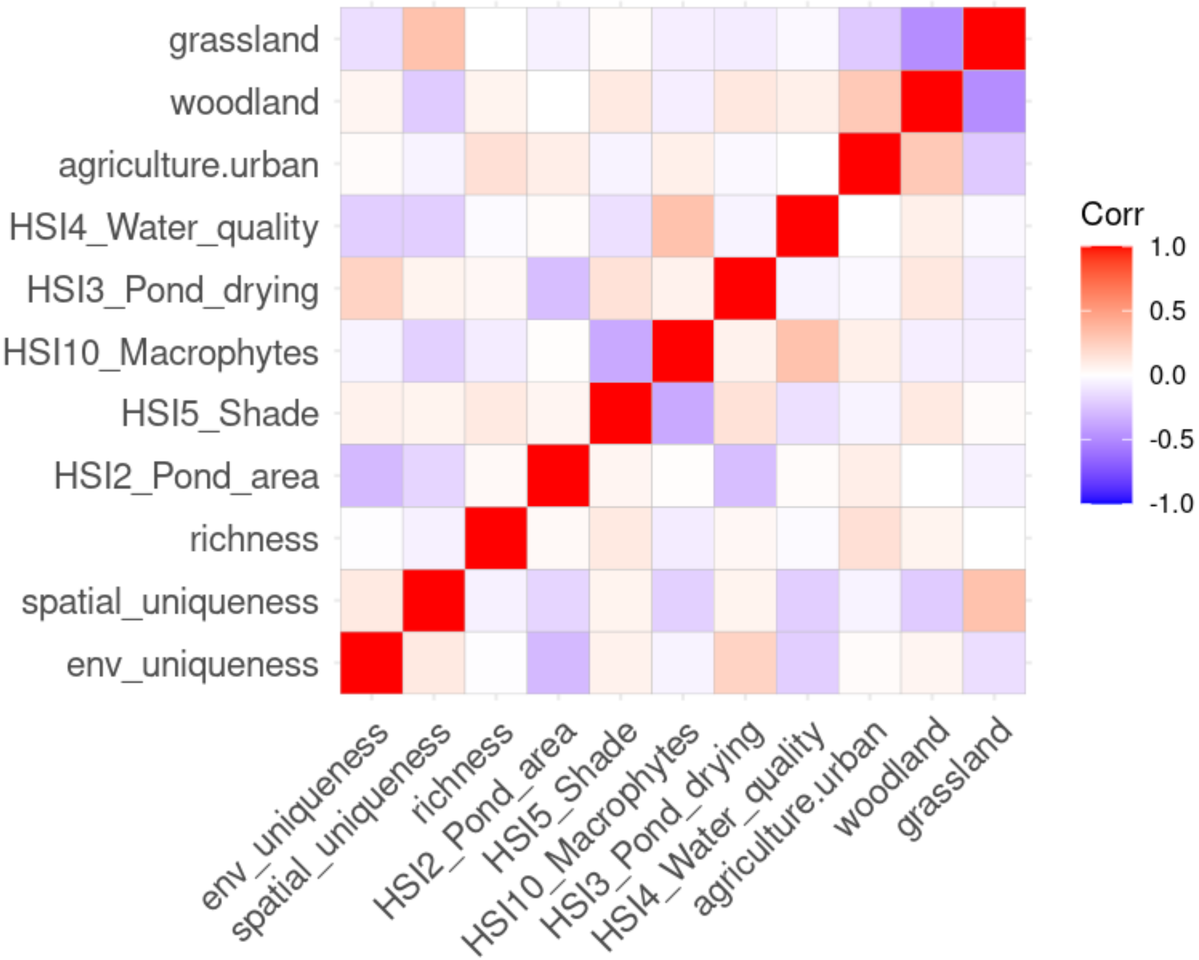

**Figure S3. Spatial distributions of the two species with high contributions of spatial autocorrelation.** A. Palmate newt *Lissotriton helveticus* (n=16 detections). B. Mandarin duck *Aix galericulata* (n=5 detections).

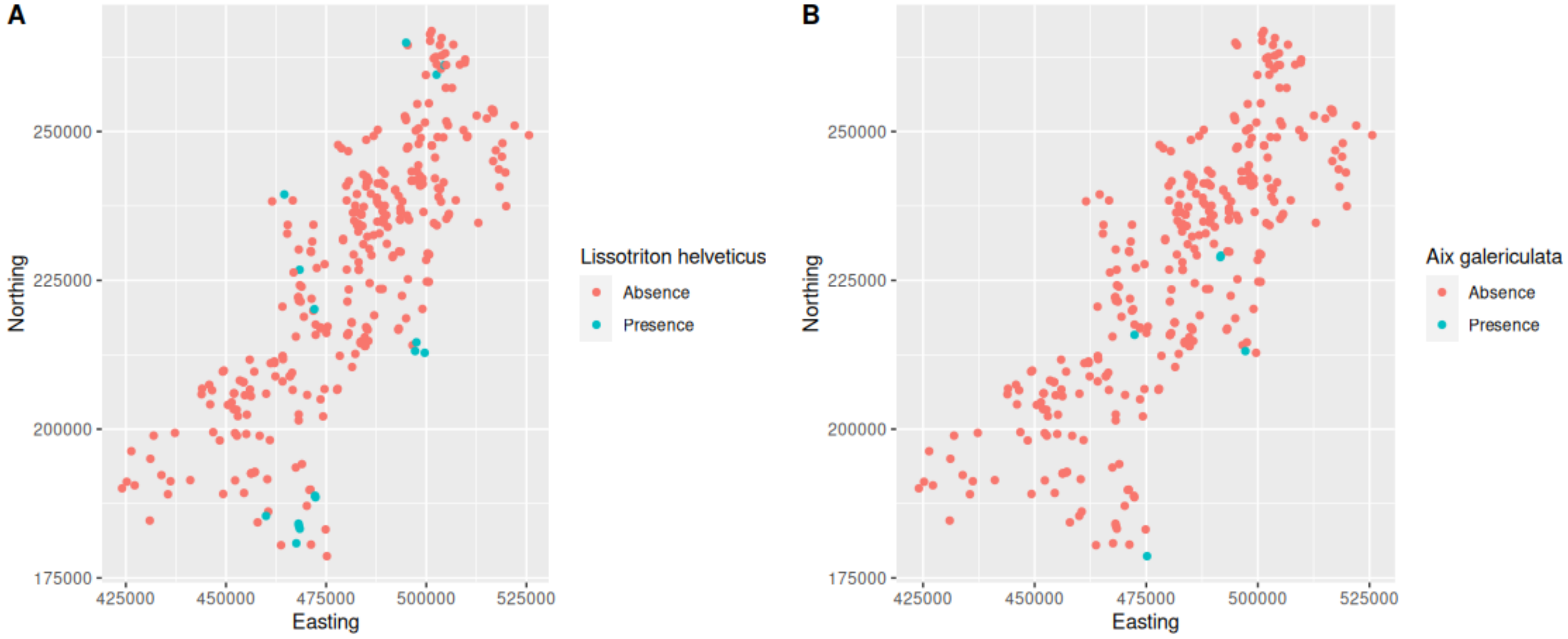

**Figure S4. Quantile regression of each model component's absolute site $R^2$ values on individual environmental covariates.** We note that if predictor variables are collinear, bivariate correlations can be spurious, and partial correlations calculated using multiple regressions should be preferred. However, the variables presented here showed practically no collinearity (Figure S4), and in the event, only one regression was significant. As this is a post-hoc demonstration of a possible analysis, no attempt is made to correct for multiple tests.

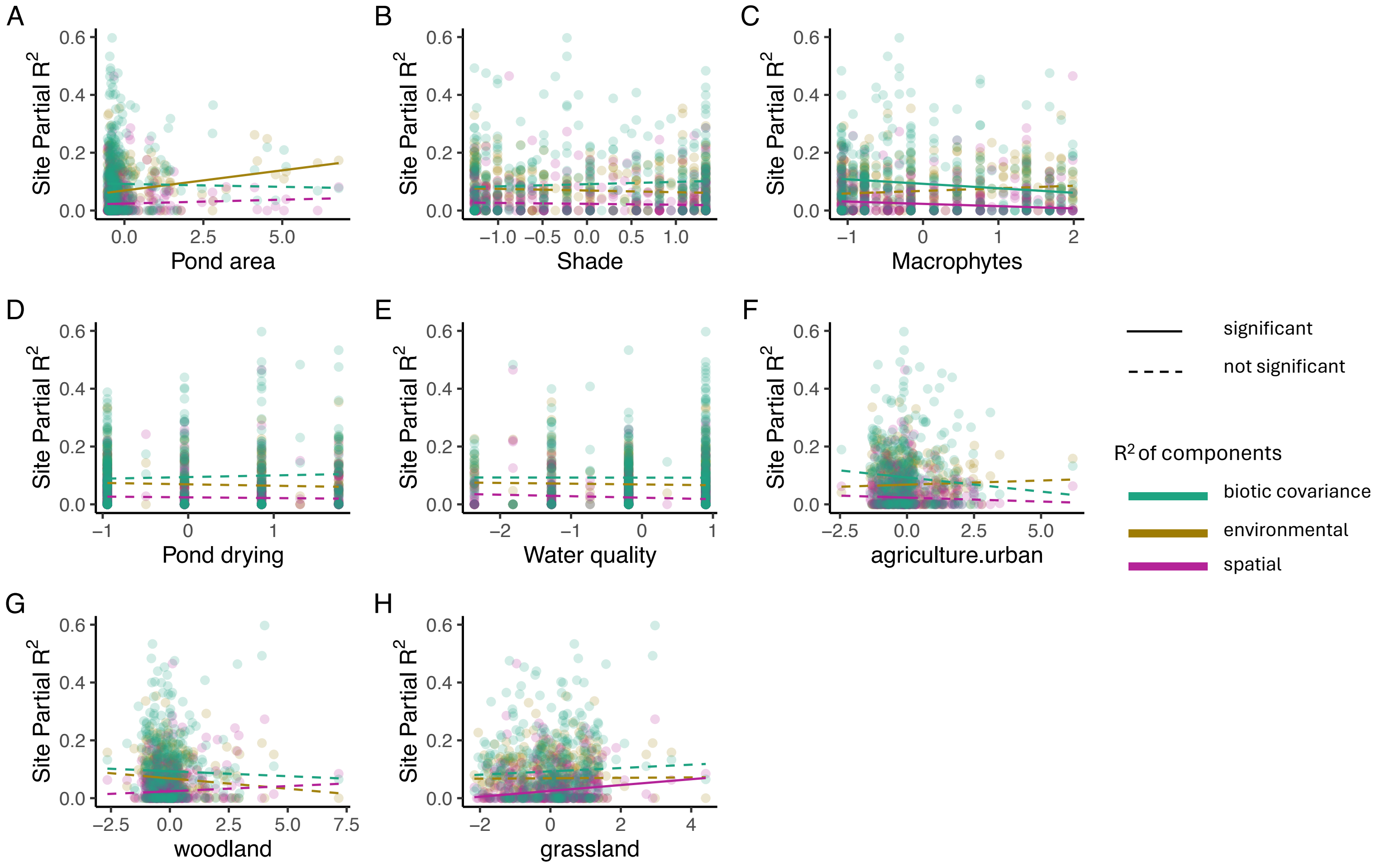

**Figure S5. Quantile regression of each model component's absolute site $R^2$ values on environmental distinctiveness, subdivided by species trait group.** For amphibians, perching birds, domestic animals, and mammals, environmental site $R^2$ significantly increases with environmental distinctiveness, which is consistent with environmental filtering acting more strongly on these taxa in environmentally distinctive sites. For more details see Main Text. As this is a *post-hoc* exploration of the robustness of the all-species result (Figure 5A), no attempt is made to correct for multiple tests.

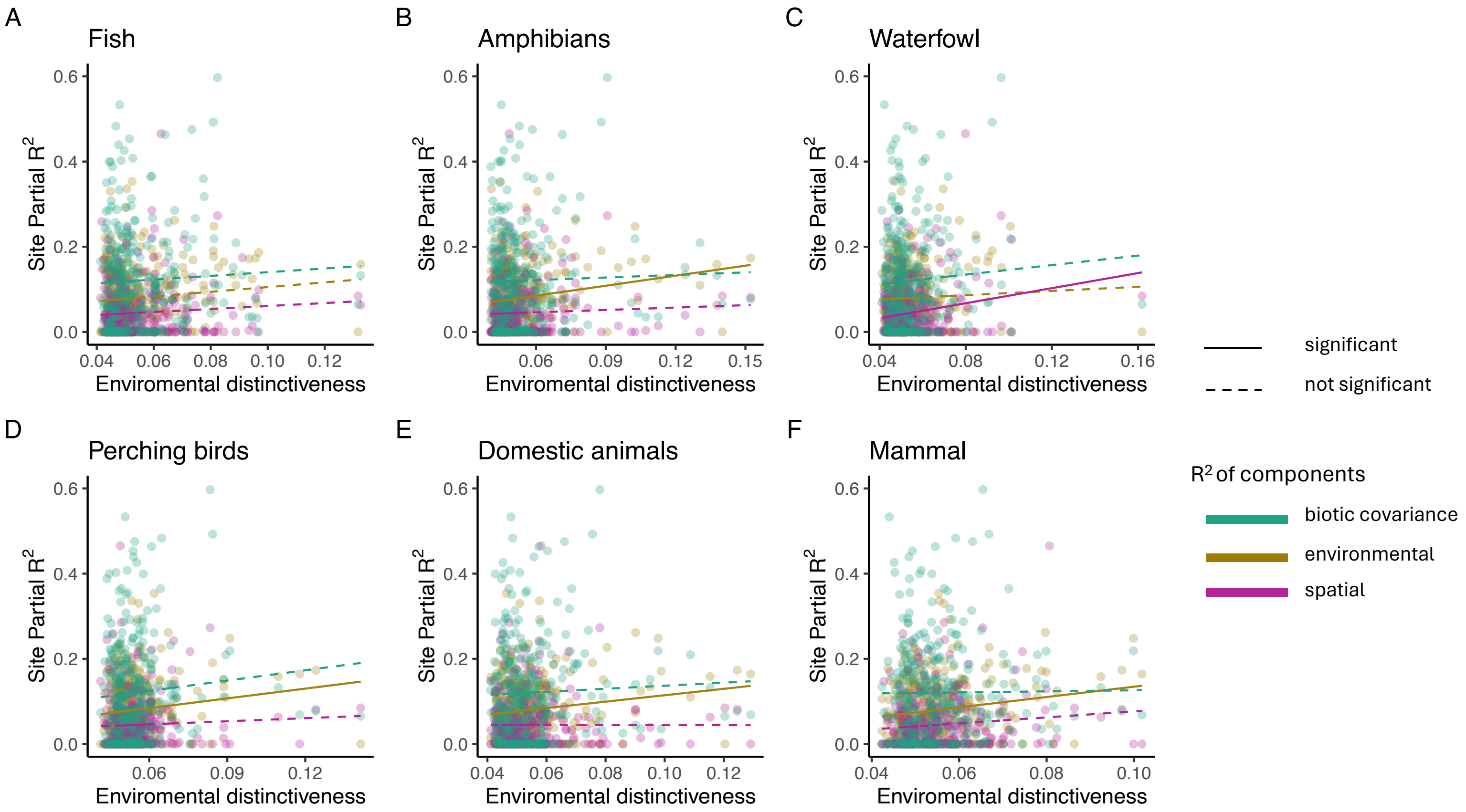
